# Interplay between phosphorylation and oligomerization tunes the conformational ensemble of SWEET transporters

**DOI:** 10.1101/2024.06.12.598708

**Authors:** Austin T. Weigle, Diwakar Shukla

## Abstract

SWEET sugar transporters are desirable biotechnological targets for improving plant growth. One engineering strategy includes modulating how SWEET transporters are regulated. Phosphorylation and oligomerization have been shown to positively regulate SWEET function, leading to increased sugar transport activity. However, constitutive phosphorylation may not be beneficial to plant health under basal conditions. Structural and mechanistic understanding of the interplay between phosphorylation and oligomerization in functional regulation of SWEETs remains limited. Using extensive molecular dynamics simulations coupled with Markov state models, we demonstrate the thermodynamic and kinetic effects of SWEET phosphorylation and oligomerization using OsSWEET2b as a model. We report that the beneficial effects of these SWEET regulatory mechanisms bias outward-facing states and improved extracellular gating, which complement published experimental findings. Our results offer molecular insights to SWEET regulation and may guide engineering strategies throughout the SWEET transport family.

## INTRODUCTION

Sugars Will Eventually be Exported Transporters (SWEETs) are bidirectional sugar uniporters that operate by transporting sugars along established concentration gradients (1). In plants, SWEETs comprise one of the major families of sugar carriers and are essential for sugar storage and distribution as it relates to plant growth, development, and defense (2). Over the years, manipulation of SWEET gene expression levels and protein transport activity has piqued interest as potential strategies to improve crop resilience and yield (3–5).

One approach to engineering SWEETs is by targeting how the protein family is regulated. Crystal structures had been elucidated, detailing how SWEETs can homo- or hetero-oligomerize to form functional complexes (6–8). Experiments have shown SWEET oligomers to be regulated in a dominant-negative fashion, where impaired function among one protomer through mutation or inhibition renders the entire complex nonfunctional (7). These results led to the belief that SWEET complexation resulted in a functional sugar-transporting pore; however, AtSWEET13 was later resolved in a monomeric form with a sugar analogue bound, indicating that SWEET monomer structures do bind sugars (8). *In silico* modeling of SWEETs resulted in a hypothesis that hetero-oligomerization could be used as a means to inhibit SWEET transport (9). Overall, it appears that the SWEET oligomerized state acts as an efficient means to ensure a rapid stop-go control of bidirectional sugar transport.

Phosphorylation is one of the most common forms of posttranslational modification in plants (10), and has been shown to increase plant sugar transport activities (11, 12). Transporter phosphorylation typically occurs along extended N- or C-terminal loops, which have evolved to be evolutionary handles for enabling cellular interactions critical to transporter function (13). A recent paper has shown that C-terminal phosphorylation of AtSWEET11 and AtSWEET12 transporters increases root:shoot ratio growth in response to drought, signifying increased sugars levels at root sink tissues in response to the abiotic stress (14). Furthermore, the same authors also showed phosphorylation to enhance oligomerization of the two transporters (14).

The current literature suggests phospho-mimetic C-terminal mutations can be introduced to crop SWEETs for engineering constitutively increased activity. Before such modification of plants is committed, the molecular details surrounding how phosphorylation and oligomerization impact SWEET transporter structure and dynamics need to be further addressed.

The function and regulation of other membrane transporter proteins also depends on an interplay between phosphorylation and oligomerization state (15). Phosphorylation can influence the form of oligomerization, as well as the resulting activity and subcellular localization of the transporter. For example, AtPIN3 auxin transporter was crystallized as a phosphorylated dimer (16). Functional characterization of PIN3 showed that without PID kinase-dependent phosphorylation, end-point PIN3-mediated efflux of indole-3-acetic acid was halved. For the nitrate transporter AtNRT1.1, a monomeric and phosphorylated state encodes low-affinity transport, whereas being dephosphorylated and dimerized encourages high-affinity transport (17–19).

Structural study of membrane proteins is notoriously challenging due to difficulties in heterologous expression and lipid reconstitution, as well as their underrepresentation in the Protein Data Bank (20). Molecular dynamics (MD) simulations serve as an attractive way to study membrane proteins due to their label-free, atomistic resolution. Modern methods in computational microscopy have greatly improved for the study of membrane proteins – especially transporters – and their posttranslational modifications (21, 22).

Using the homotrimer crystal structure of OsSWEET2b (PDB ID: 5CTG) (7), we employ large scale MD simulations of monomeric and oligomeric SWEET proteins under different phosphorylation states. We report how both forms of SWEET regulation directly impact transporter gating and the implications of our results in potential SWEET engineering strategies.

## RESULTS

### Identifying putative phosphorylation sites for OsSWEET2b

OsSWEET2b was selected for study because of the availability of its oligomeric crystal structure (7). The structure was resolved as a homotrimer of three inward-facing (IF) protomers (**Figure 1**). Using bioinformatic software, putative OsSWEET2b phosphorylation sites were predicted (**Table S1**). Probable C-terminal phosphorylation sites included Tyr214, Ser215, Ser223, and Ser224. While the OsSWEET2b C-terminal tail is shorter than other SWEETs (17 residues rather than 20–40+ residues), it presented a similar predicted phosphorylation pattern compared to those known for other SWEETs. For example, two AtSWEET11 and AtSWEET12 phosphorylation sites were found to be Ser237 and Ser248 (14). Likewise, OsSWEET2b predicted phosphosites also total two general regions on the tail separated by an ∼8 residue spacer.

**Figure 1.**
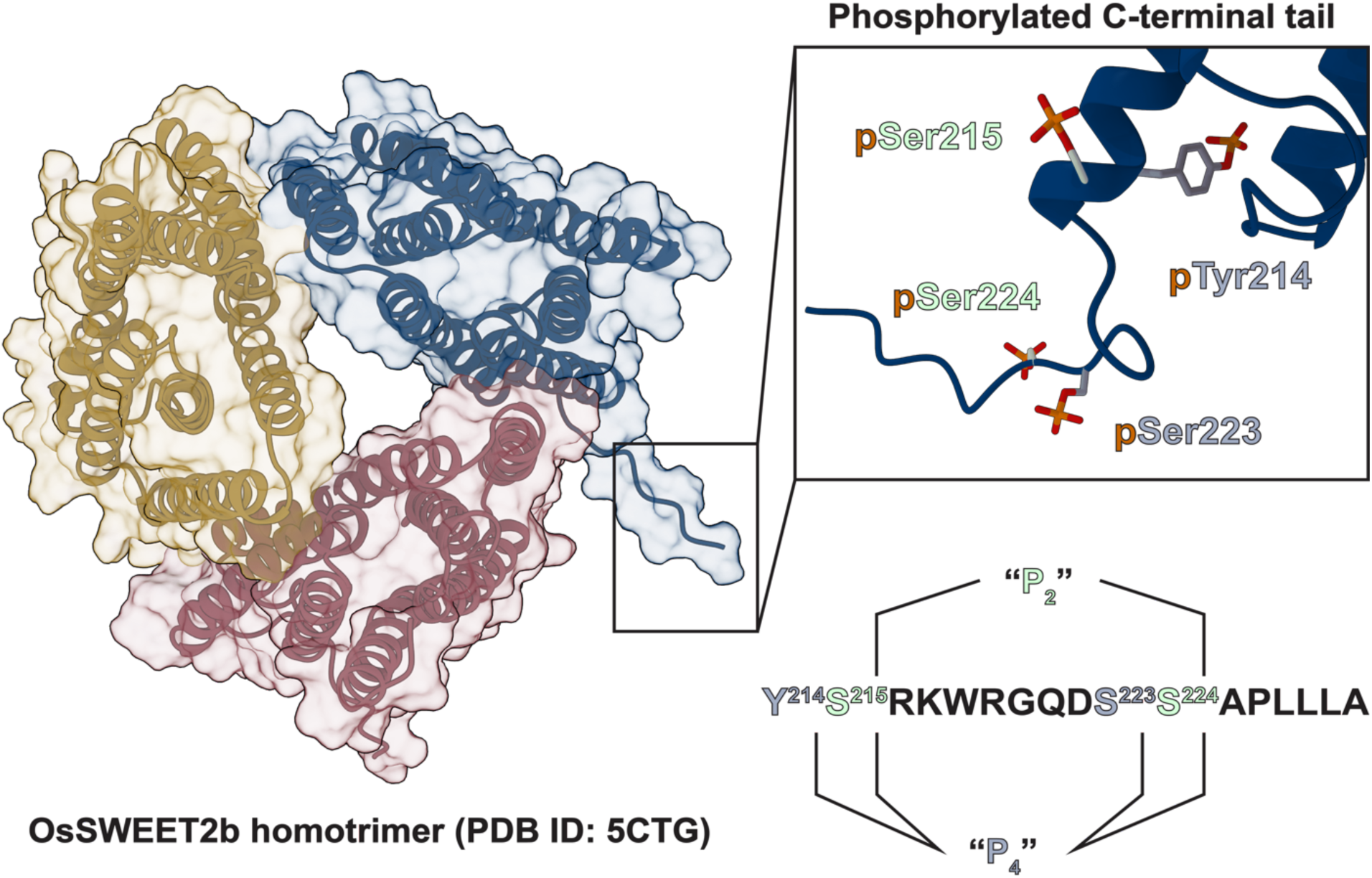
Overview of predicted C-terminal phosphorylation sites with respect to OsSWEET2b homotrimer structure. Simulations were performed using either the “P_2_” (pSer215, pSer224) or the “P_4_” phosphorylation pattern (pTyr214, pSer215, pSer223, pSer224). Phosphorylation sites were selected based off bioinformatic predictions and comparison to experimentally validated patterning for AtSWEET11 and AtSWEET12.

To account for the possible combinatorics in phosphorylation patterns, we modeled two types of phosphorylation systems – pSer215/pSer224 to match the AtSWEET11/12 patterns (“P_2_”), and pTyr214/Ser215/Ser223/Ser224 to consider an extreme case of phosphorylation-dependent effects (“P_4_”). Each of these patterns, along with a non-phosphorylated variant, was modeled using both monomer and trimer OsSWEET2b structures. Resulting data reflecting phosphorylation and oligomerization status are described as p_0_OsSWEET2b^M^, p_2_OsSWEET2b^M^ and p_4_OsSWEET2b^M^ for the monomer; and p_0_OsSWEET2b^T^, p_2_OsSWEET2b^T^ and p_4_OsSWEET2b^T^ for the trimer.

### Phosphorylation of monomeric OsSWEET2b promotes outward-open states

In agreement with past simulation studies, OsSWEET2b was found to have sampled hourglass (HG), inward-facing (IF), occluded (OC), and outward-facing (OF) conformations (23–27), meaning our OsSWEET2b monomer simulations were able to capture the entire conformational cycle regardless of phosphorylation state (**Figure 2A**). Free energy values for monomer OsSWEET2b metastable states across each phosphorylation pattern are summarized in **Figure 3**.

**Figure 2.**
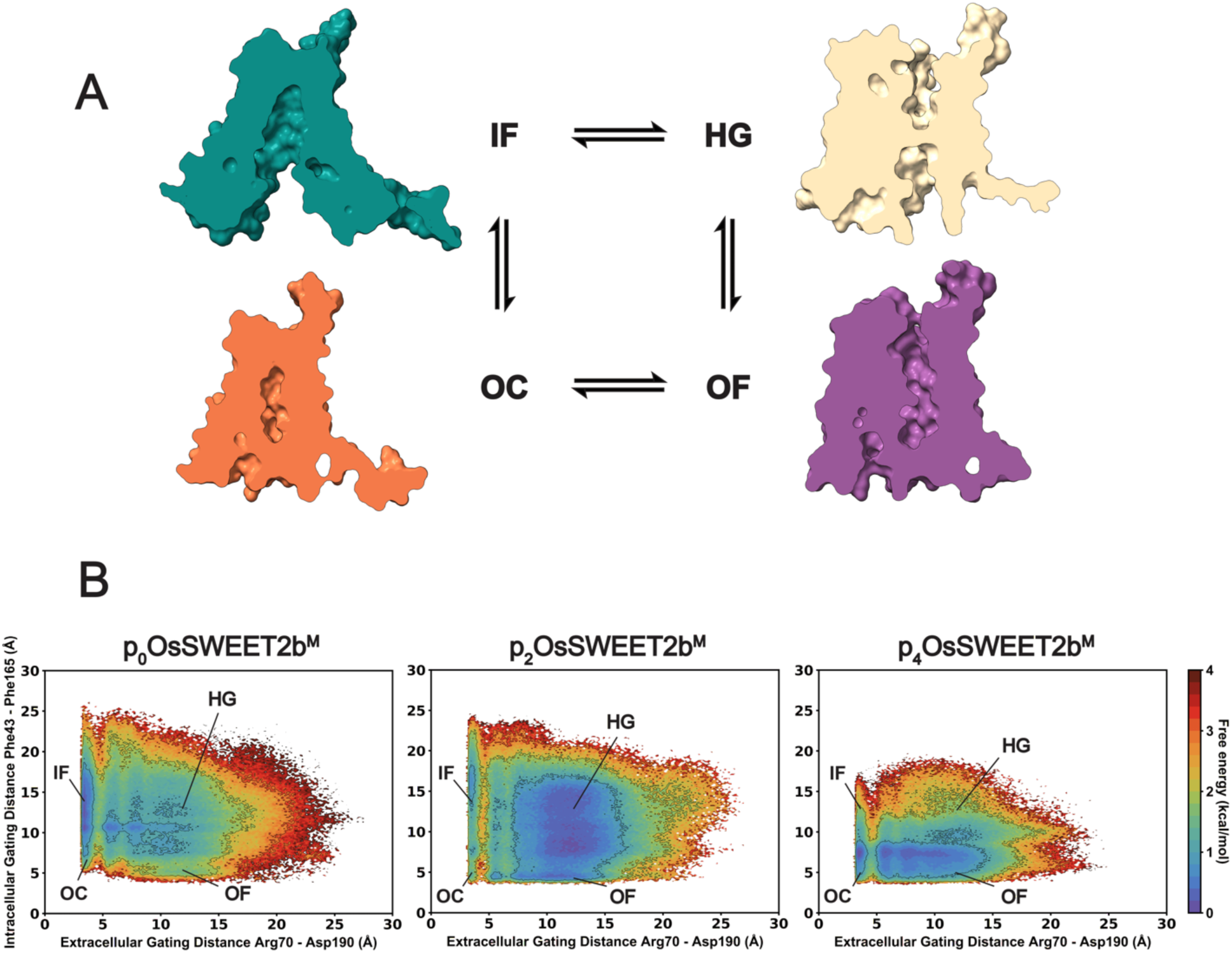
Conformational sampling by the OsSWEET2b monomer in response to different phosphorylation states. (**A**) Representative structures for the inward-facing (IF), occluded (OC), hourglass (HG), and outward-facing (OF) conformational states sampled by monomeric OsSWEET2b throughout *apo* alternate access. (**B**) Gating landscapes comparing the apertures for OsSWEET2b extracellular gate versus the intracellular gate under no (*left*; 51.94 µs aggregate simulation), P_2_ (*middle*; 55.58 µs aggregate simulation), and P_4_ (*right*; 54.04 µs aggregate simulation) phosphorylation patterning. Gating distances are plotted in angstroms (Å) with free energy reported in kcal/mol.

**Figure 3.**
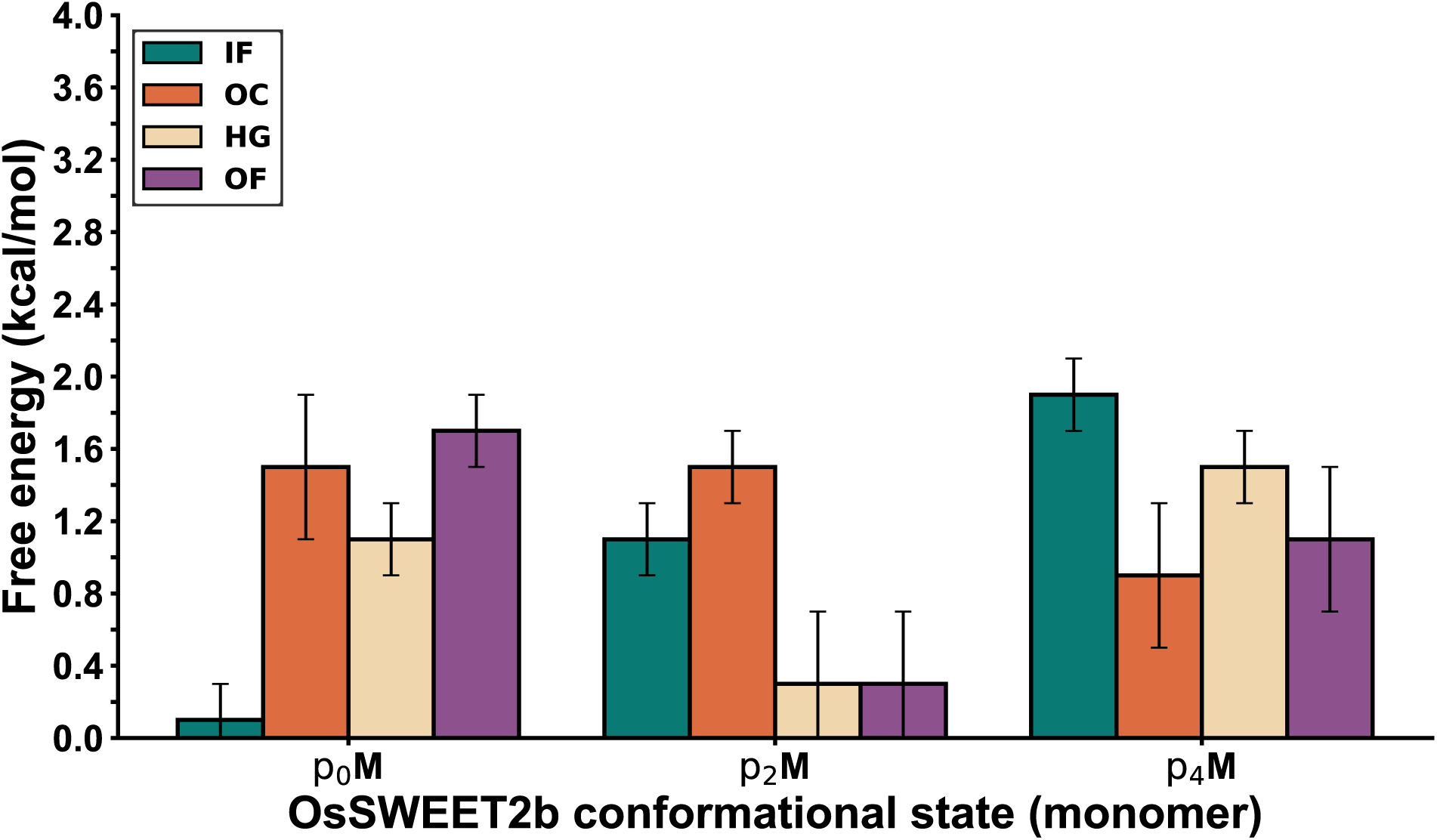
Gating state thermodynamics for monomeric OsSWEET2b. Free energy reported in kcal/mol based off landscapes shown in Figure 2B. Error bars are shown based off bootstrapped landscapes in **Figure S5**.

Without phosphorylation, p_0_OsSWEET2b^M^ is most likely to sample an IF state at a relative free energy of 0.2–0.6 ± 0.2 kcal/mol (**Figure 2B**, *left*). The most thermodynamically favorable conformational transitions occur through an intermediate IF-OC state, where the intracellular gating distances approximates 10 Å and the extracellular gating distances remains around 5 Å. This result represents a population shift from our previous study where p_0_OsSWEET2b^M^ was simulated without the C-terminus (26), indicating that inclusion of the tail redirects the free energy landscape away from IF-like intermediate states. Through this transition, p_0_OsSWEET2b^M^ samples an OF state at a relative free energy of 1.6–2.0 ± 0.2 kcal/mol. Without phosphorylation, p_0_OsSWEET2b^M^ intermediate states are likely to reflect this ∼1.4 kcal/mol energy difference by preferring a more IF-like conformation, where sampling HG or extended IF states become as probable as sampling an OF state.

Following the AtSWEET11/12 phosphorylation pattern, p_2_OsSWEET2b^M^ of an IF state becomes ∼0.5 kcal/mol less stable (**Figure 2B**, *middle*). Meanwhile, a large free energy barrier of 2.4–2.6 ± 0.2 kcal/mol now separates strictly IF states from opening the extracellular gate. After crossing this barrier, the same IF-OC intermediate state seen in the p_0_OsSWEET2b^M^ landscape is ∼0.8 kcal/mol less stable, likely redirecting the p_2_OsSWEET2b^M^ ensemble preference towards the relatively flat basins of HG or OF states (both 0.2–0.4 ± 0.4 kcal/mol).

The extreme case of p_4_OsSWEET2b^M^ follows similar trends as p_2_OsSWEET2b^M^, except sampling the IF state becomes more difficult (**Figure 2B**, *right*). IF- or HG-IF-like states typically incur free energy costs of 2.2 ± 0.2 kcal/mol, making them the least favorable functional states out of all three phospho-ensembles. Furthermore, an intracellular gate distance greater than ∼13 Å becomes very difficult to achieve, meaning p_4_OsSWEET2b^M^ would likely remain in either an OC, OC-OF, or OF state. These results suggest that p_4_OsSWEET2b^M^ would not be able to efficiently commit alternate access and function within plants.

Our simulation results suggest a p_2_OsSWEET2 pattern, which corroborates with the experimentally validated “2P” patterns seen for AtSWEET11/12 (14).

### Oligomerization rescues inward-open states and maintains independent gating between protomers

Although SWEET proteins can be functional as monomers, OsSWEET2b was crystallized as a homotrimer (7). Each of the protomers within the OsSWEET2b trimer was resolved in the same IF conformation. From this result, the crystallographers hypothesized that each protomer could function as independent sugar transport routes. However, C-terminal phosphorylation could modulate these proposed allosteric coupling between protomers. Thus, we proceeded with simulations of OsSWEET2b trimer across the different phosphorylation states (**Figure 4**). As for the monomer results, the free energy values for metastable states seen for each protomer are shown in **Figure 5**.

**Figure 4.**
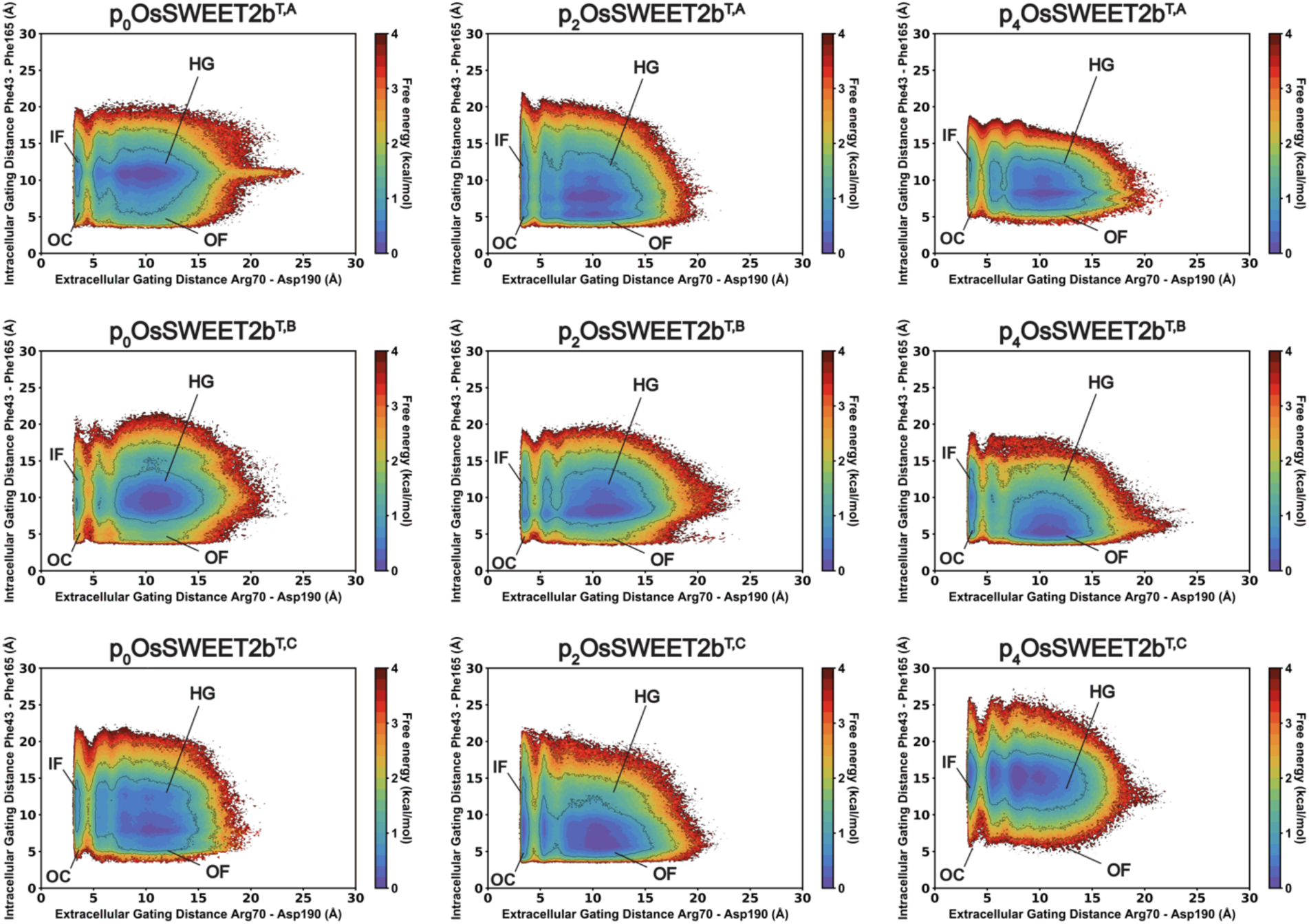
Combined effect of phosphorylation and oligomerization on OsSWEET2b. Gating landscapes compare shown for no (*left column*; 588.62 µs aggregate simulation), P_2_ (*middle column*; 591.57 µs aggregate simulation), and P_4_ (*right column*; 593.40 µs aggregate simulation) phosphorylation patterning. Individual protomers are shown for chains A (*top row*), B (*middle row*), and C (*bottom row*). Gating distances are plotted in angstroms (Å) with free energy reported in kcal/mol.

**Figure 5.**
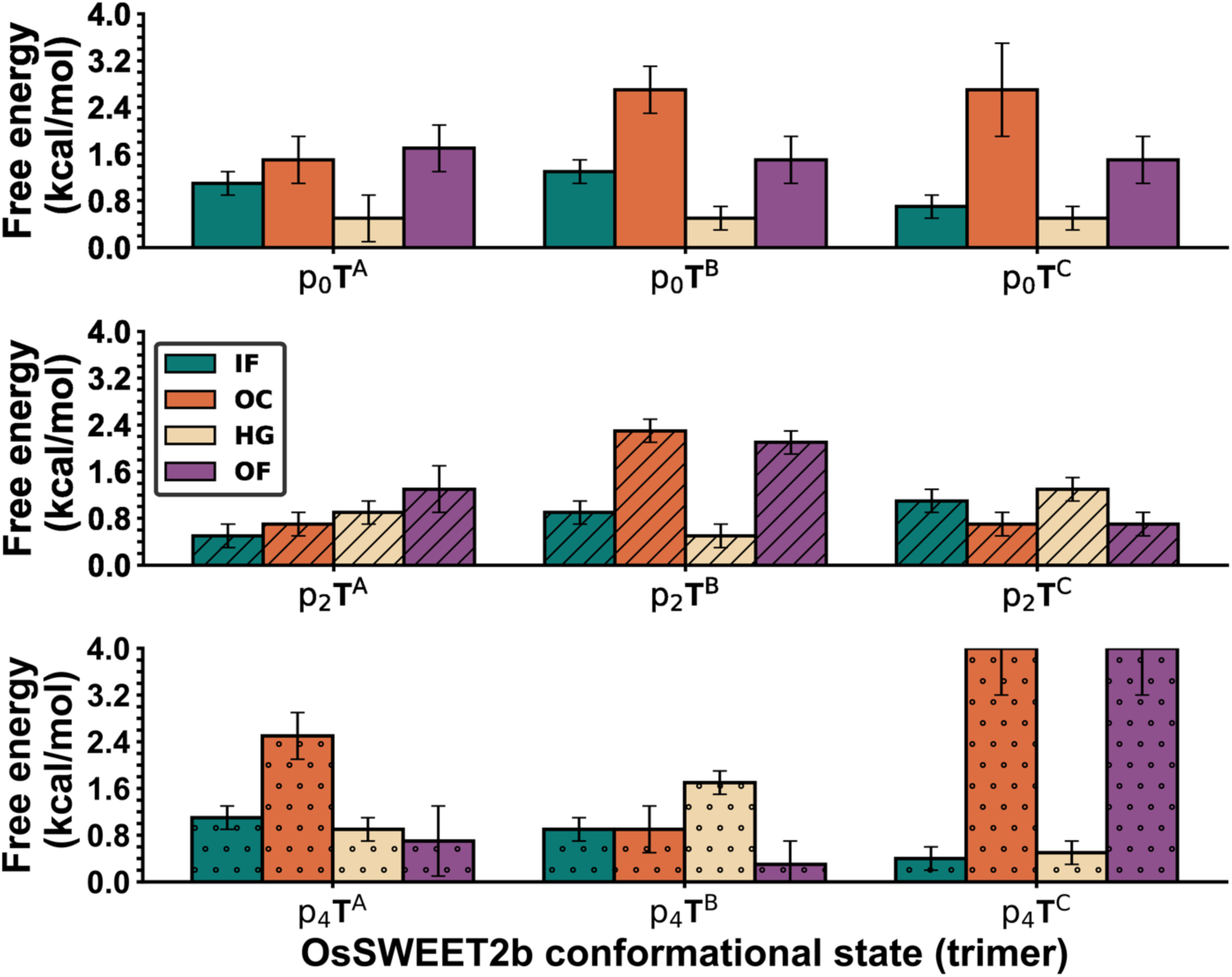
Gating state thermodynamics for trimeric OsSWEET2b protomers. Phosphorylation patterns by row: (*Top row*) P_0_, (*middle row*) P_2_, and (*bottom row*) P_4_. Free energy reported in kcal/mol based off landscapes shown in Figure 4. Error bars are shown based off bootstrapped landscapes in **Figure S6**.

Incorporation of OsSWEET2b into a trimeric complex enforces strict separation of metastable states while offering specific stabilization effects with respect to gating. Compared to monomer simulations, IF states are stabilized within the trimer. Just like for the monomer simulations, each OsSWEET2b protomer can still capture the entire conformational cycle, regardless of phosphorylation status.

Without phosphorylation, each of the p_0_OsSWEET2b^T^ protomers present an effective energy barrier between IF and OC-IF or OF-like states anywhere between 1.6–2.6 ± 0.2 kcal/mol (**Figure 4**, *left*). The p_2_OsSWEET2b^T^ and p_4_OsSWEET2b^T^ variants present similar barriers towards an IF-to-OF transitions. p_0_OsSWEET2b^T^ IF states are destabilized by 0.2–1.2 kcal/mol when compared against p_0_OsSWEET2b^M^ sampling (Figure 2, *left*). Interestingly, p_2_OsSWEET2b^T^ IF states are 0.2–0.4 kcal/mol more stable than when sampled as p_2_OsSWEET2b^M^ (**Figures 2 and 4**, *middle*). IF stabilization for p_4_OsSWEET2b^T^ is even more pronounced, lowering the free energy from 2.2 ± 0.2 kcal/mol as seen for p_4_OsSWEET2b^M^ down to a range of 0.4–1.0 ± 0.2 kcal/mol (**Figures 2 and 4**, *right*).

Besides rescuing IF states, the phosphorylation of trimeric OsSWEET2b further maintains OF-like stability seen from monomer simulations. OF states are still destabilized in favor of more IF-like or HG-IF states for p_0_OsSWEET2b^T^, except in the event of protomer p_0_OsSWEET2b^T,A^. Agreement between p_2_OsSWEET2b^M^ and p_2_OsSWEET2b^T^ is maintained by greater OF and HG stabilization at a sub-1.0 kcal/mol level, although p_2_OsSWEET2b^T,B^ samples strictly OF states at a free energy of 1.6–2.0 ± 0.2 kcal/mol (**Figure 4**). Protomers of p_4_OsSWEET2b^T^ still stabilize HG-OF or OF-like states relative to p_0_OsSWEET2b^T^ (**Figure 4**, *right*).

Teasing apart the importance of (functional) differences between complex-participating protomer gating is complicated. Literature suggests that oligomerization stabilizes SWEET transporters and enables increased activity (6, 7, 14). Overall, our free energy landscapes suggest that the overstabilization of OF-like states in p_2 or 4_OsSWEET2b^M^ is counterbalanced by a general stabilization of IF states as well. Without phosphorylation a more marked imbalance of IF versus OF sampling remains. Comparing each trimer gating landscape, a trend emerges where two out of the three protomers appear to sample closely similar distributions, or at least manage similar extents of IF:OF stabilization (e.g., p_0_OsSWEET2b^T,A+B^ or p_2_OsSWEET2b^T,A+C^).

The idea of inter-protomer coupling has been mentioned in SWEET structural biology, where OsSWEET2b protomers were predicted to gate independently (7, 8). To test this, we monitored the time series evolution of protomer-specific gating using representative trimer trajectories. One of the three protomers was fixed as either an IF-like or OF-like state, while the remaining two could vary. Our analyses did not reveal any conclusive coupling. Instead, we found that each of the protomers present within the homotrimer could simultaneously sample independent conformations (**Figure 6**). One such trajectory – p_4_OsSWEET2b^T^ #452 – is represented in **Figure 6** because each respective protomer samples either the IF, OF, or HG state. We conclude that protomer dynamics are independent, which extends to the possibility of sampling similar metastable states at the same time. Alternatively, with respect to their dominant-negative regulation (7), there also exists the possibility where the conformational state of some SWEET protomers may weakly restrict the sampling performed by others (**Figure 4**).

**Figure 6.**
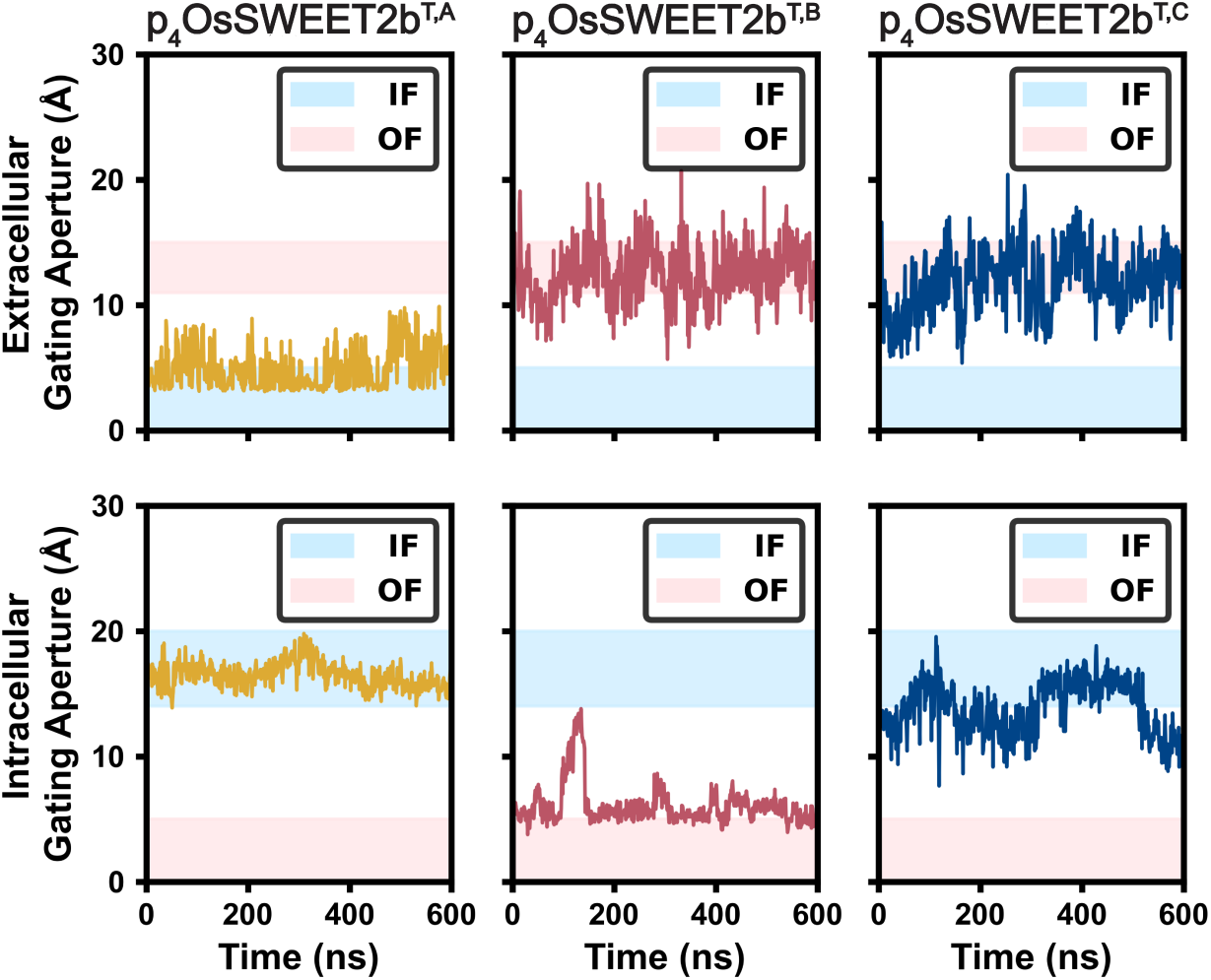
Independent gating observed amongst OsSWEET2b protomers. Extracellular gating aperture (*top row*) is compared against intracellular gating aperture (*bottom row*) for an example timer trajectory where distinct metastable states were simultaneously observed per protomer (*Chain A*, *left*; *Chain B, middle; Chain C, right*). If both time-series sample the blue-highlighted aperture, the protomer is in the IF state. If both time-series sample the pink-highlighted aperture, the protomer is in the OF state. If opposing highlighted apertures are sampled simultaneously, the protomer is in the OC or HG state.

### Phosphorylation and oligomerization kinetically regulate OsSWEET2b gating transitions

Our thermodynamic analyses suggest that phosphorylated and oligomerized OsSWEET2b energetically prefer different conformational states. Therefore, these post-translational modifications must likely also impact gating rates of transition.

Because AtSWEET11/12 phosphorylation and oligomerization contribute to improved root growth under drought stress (14), that would mean root sugar levels are somehow improved. Source-to-sink sugar transport can be improved *in planta* by (1) making sugar sink-directed transport rates faster or (2) making sugar source-directed transport rates rate-determining. Since soure:sink relationships can change between plant tissues, the direction of sugar transport cannot be consistently described as import or export. With respect to our simulations, we can only describe whether OsSWEET2b faces the outside of the cell or the inside of the cell. Directional transport can then be approximated by calculating the rates of OF-to-IF conformational transition and the rates of IF-to-OF conformational transition (**Figure 7**). The exact values and statistics for these computationally predicted rates are provided in **Table S2**.

**Figure 7.**
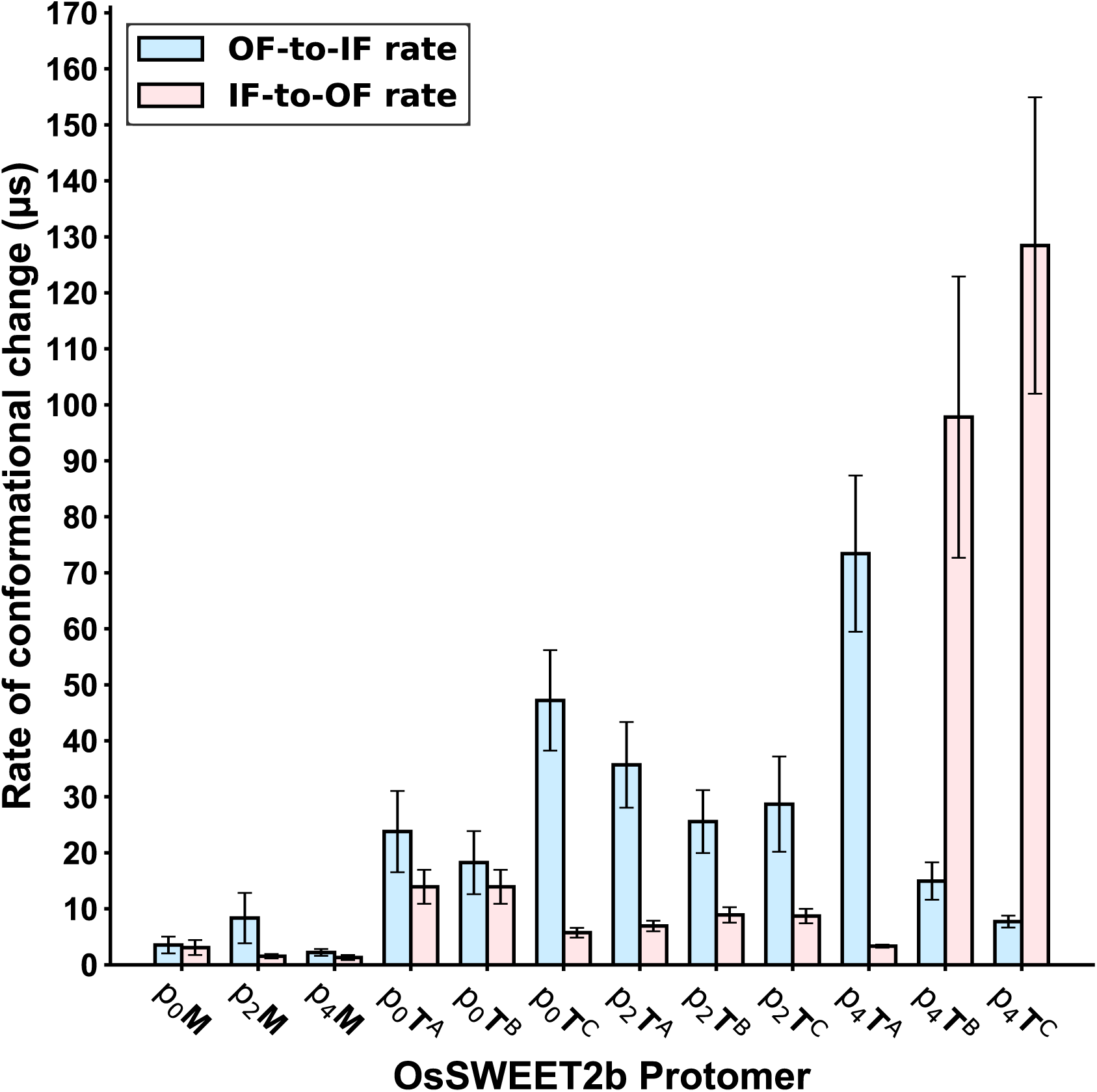
Rate calculations for OF-to-IF or IF-to-OF OsSWEET2b conformational change.

For OsSWEET2b^M^, the results show IF-to-OF rates as faster than OF-to-IF rates for all cases. The two rates for p_0_OsSWEET2b^M^ are within error at 3.0789 ± 1.3474 µs and 3.5287 ± 1.500 µs, respectively. Unphosphorylated trimer IF-to-OF rates again remain faster, with a significant difference in rates for p_0_OsSWEET2b^T,C^. Comparing pOsSWEET2b^T^ to p_0_OsSWEET2b^M^, trimer rates for OF-to-IF are ∼15–35 µs slower. But the trimer IF-to-OF rates are only ∼3–10 µs slower than monomer rates. Without phosphorylation, it appears that OsSWEET2b IF-to-OF rates are simply faster than those for OF-to-IF.

However, when combining phosphorylation and oligomerization for p_2_OsSWEET2b^T^, IF-to-OF rates are not only faster, but OF-to-IF rates are roughly three to five times slower (indicated by non-overlapping error bars and extreme statistical significance; **Figure 7**, **Table S2**). Given that the P_2_ putative phosphorylation pattern is more likely to exist, we expect these results to hold greater value than the p_4_OsSWEET2b^T^ rates, which show a reversal in favor of faster OF-to-IF rates instead (**Figure 7**).

Overall, both phosphorylation and oligomerization alone promote faster rates of transition from an IF to OF states. Together, combined phosphorylation and oligomerization make gating transitions from OF to IF states rate-determining. Without phosphorylation, IF-to-OF and OF-to-IF transition rates are close enough for OsSWEET2b^T^ to accommodate sugar transport flowing in any direction. On the contrary, OsSWEET2b trimer phosphorylation appears to act as a key regulatory mechanism in maintaining when and how directional sugar transport is preferred.

## DISCUSSION

SWEETs and their sugar-transporting functions are critical for general plant physiology. Therefore, these transporters become important targets for improving food security and crop yield. Developing effective strategies for engineering SWEET function will become a paramount research question as food demands increase alongside the human population. Targeting the mechanisms regulating SWEET function could help serve as useful and orthogonal means to currently employed engineering tactics, such as increasing SWEET transcription at the promoter level. Still, engineering SWEET regulation cannot proceed without better coverage of the molecular details. We provide this information using MD simulations of monomeric and trimeric OsSWEET2b under varying levels of phosphorylation.

Increasing phosphorylation led to a greater preference towards OF-like states in monomeric OsSWEET2b. When incorporated into trimers, the effects of phosphorylation and oligomerization counterbalanced one another, leading to stabilized IF-like states but maintained OF or HG stabilization when compared to phosphorylated monomers. Conversely, non-phosphorylated OsSWEET2b experienced equivalent difficulties in sampling an OF-like state regardless of oligomerization status. Our thermodynamics and kinetic analyses lead us to suggest that the combined effect of phosphorylation and oligomerization results in the enhancement of outward-facing conformational changes. These findings agree with *in planta* experiments that showed phosphorylated and oligomerized AtSWEET11/12 increased sugar export towards root tissues during drought conditions (14).

Another routine analysis when modeling phosphorylated proteins is looking at hydrogen bond analyses (28). Because phosphorylation was reported to increase C-terminal contacts with SWEET cytosolic loops from yeast two-hybrid experiments (14), we evaluated whether hydrogen bonding interactions could reflect this finding. Our hydrogen bonding analyses found C-terminus–cytosolic loop interaction lifetimes to increase after phosphorylation (**Tables S3 and S4**), providing confidence that the structures we have obtained and analyzed are capturing the same phenomenon seen from AtSWEET11 and AtSWEET12 experiments (14). Thus, our results appear to confirm a generalized behavior across SWEET transporters with C-terminal loops.

Our simulation results confirm the previous hypothesis that each of the OsSWEET2b protomers can function independently when assembled into a trimeric complex (7). This finding also agrees with the notion that SWEET transporters can hetero-oligomerize, as each protomer would still function independently (6). SWEET transporters within the same clades can recognize similar substrates even without sharing subcellular localization (29). A benefit to having independent protomers is the greater availability of sugar apparent transport routes. On the other hand, the potential for each protomer to be in a different state means that mechanisms for regulating directional transport becomes less efficient. Our kinetic rate analyses suggest that phosphorylation is the key biophysical determinant of directional sugar transport by oligomerized SWEETs. Since sugar transport is governed by source-to-sink relationships, the ideal scenario is for each protomer to share the same conformation when being directionally controlled upon phosphorylation.

The seemingly universal response of SWEETs to phosphorylation is interesting, as SWEETs present different spatiotemporal expression patterns across different plant tissues and developmental stages (30). For instance, OsSWEET2b is expressed in the tonoplast and is responsible for regulating sugar flux to and from the vacuole. OsSWEET2b phosphorylation would likely lead to a sugar-starved vacuole, as IF-to-OF transitions would become more favorable. Loss of vacuolar AtSWEET2 has been shown to offer resistance towards pathogenic root-colonizing fungi (31), suggesting that increasing sugar sequestration away from the vacuole could be harmful to plant health. Alternatively, increasing basal source-to-sink sugar export could also improve relations between plants and mycorrhizae symbionts (5). A delicate physiological balance exists between reserving and dispensing sugars, and the functional redundancy of plant transporters expressed to manage these processes (4).

Although improved sink-directed sugar transport by SWEETs has proven useful under (a)biotic stress, there are also drawbacks to this approach. Overexpression of some Clade III SWEETs derive impaired growth phenotypes due to deficient sugar transport; this is done at the expense of increased pathogen resistance (e.g., StSWEET11, AtSWEET11, OsSWEET11, AtSWEET12, OsSWEET14, AtSWEET14, AtSWEET15; reviewed in Ref. (4)). Importantly, this is not the case for all Clade III SWEET transporters, nor for SWEETs belonging to different clades. It has been suggested that increasing phosphorylation and oligomerization could dynamically increase SWEET activity (4). Yet mutations that lead to constitutive phosphorylation may also lead to similar harmful effects akin to SWEET gene overexpression under basal conditions. This is likely because of the predicted increased stringency on directional sugar transport upon phosphorylation.

A potential compromise could be to perform a mutagenesis scanning along the cytosolic loops with a phospho-mimetic mutation along the C-terminus. Our hydrogen bonding analyses show interactions between the C-terminal tail and cytosolic loops increase upon phosphorylation (**Table S3 and S4**). Engineering improved SWEET transporters may require a phosphorylation-compatible mutation that preserves biological feedback mechanisms to (a)biotic stressors without forcing constitutive activity.

In conclusion, our observations strengthen experimentally proposed hypotheses about the physiological implications of SWEET regulation by phosphorylation and oligomerization. In turn, these results offer potential strategies for engineering these important plant membrane transporters.

## EXPERIMENTAL PROCEDURES

### Prediction of phosphorylation sites

The OsSWEET2b amino acid fasta sequence (UNIPROT ID: Q5N8J1) was used to query putative phosphorylation sites using the following tools: GPS-6.0, NetPhos-3.1, and MusiteDeep (32–34). Attention was paid towards the C-terminal tail. Consensus of scores for different suggested kinases indicated that residue sites should be recognizable by generalized kinase domains. Comparison to *Arabidopsis* SWEET transporters suggested a two-site (P_2_) phosphorylation pattern (14), whereas a four-site (P_4_) pattern could suggest extreme consequences of phosphorylation. A P_2_ pattern was decided to comprise Ser215/Ser224, whereas the P_4_ pattern comprised Tyr214/Ser215/Ser223/Ser224.

### Modeling of C-terminal tail

The C-terminal tail was extended for each starting OsSWEET2b PDB file using the cyclic coordinate descent approach in Rosetta software using the cyclic coordinate descent protocol (Ty214 – Ala230; 17 total residues) (35, 36). Given how the C-termini in the OsSWEET2b homotrimer crystal structure (PDB ID: 5CTG) lack secondary structure (7), the resulting Rosetta C-termini output were linearized using MODELLER (37). The DOPE score was maximized across 100 structures to determine an appropriate tail conformation for simulations. This protocol was repeated for each OsSWEET2b monomer starting conformation, as well as each protomer within the homotrimer crystal structure.

### System Construction

PDB files with modeled C-termini were supplied to CHARMM-GUI for system construction, where missing hydrogens were also added (38). Such PDB files include an IF, OF, OC, and HG conformation for monomeric OsSWEET2b (24, 26), the OsSWEET2b homotrimer, and each of their respective phosphorylated variants. The CHARMM36m force fieId was used for parameterization (39). A realistic plant plasma membrane bilayer model composition was used for simulation of OsSWEET2b proteins (26). Maximum complexity recipes (26) were used at sizes of 256 and 512 total lipids for monomer and trimer system construction, respectively. A distinct membrane packing was initialized for each of the input PDB states to help prevent unwanted lipid ensemble properties from impeding conformational sampling during simulations (26, 40, 41). The CHARMM default TIP3P water was used for solvation (42). Neutralizing K^+^ and Cl^-^ ions were added to a concentration of 0.15 M. In accordance with the intracellular pH of plant cells as ∼7.5, the SP2 patch was used for phosphorylation of Ser215/Ser223/Ser224, while the TP2 patch was used for Tyr214. Hydrogen mass repartitioning was used to enable a 4 fs timestep (43).

### Molecular dynamics simulation parameters

OpenMM 7.7.0 was used to perform classical molecular dynamics simulations locally and on Folding@Home (44–46). Energy minimization, NVT heating, NPT heating and equilibrations were performed according to standardized best practices (47, 48). Local energy minimization was performed for 50000 steps. NVT heating was then performed from 0K to 310K for 1 ns. Subsequent NPT simulation was performed for 3 ns. Next, a 50 ns NPT equilibration run was performed as a “HOLD” step (26, 40). Each of these above preparatory simulations were performed using protein backbone harmonic restraints at a strength of 5 kcal/mol/Å^2^. Backbone restraints were then lifted for an additional 50 ns equilibration run. Lastly, a 100 ns production run was performed prior to adaptive sampling. The Particle-Mesh-Ewald method was used for treatment of electrostatic forces (49), where the nonbonded cutoff was selected as 12 Å in agreement with the CHARMM force field. Hbond constraints and rigid water settings were turned on for all simulations. A Langevin thermostat was used for maintaining simulation temperature with a Langevin friction coefficient of 2.8284 ps^-1^. A Monte Carlo Membrane barostat with isotropic pressure scaling was used for maintaining a pressure of 1.0 atm. Membrane surface tension was maintained at 200•bar•nanometer.

### Adaptive sampling regime

Adaptive sampling was used to enhance sampling by selecting seed states from short unbiased trajectories (50–54). Adaptive sampling is performed progressively in rounds, where any frame from a simulation completed in one round can be used as a starting point, or seed, to begin a simulation in the next round. Seed states are selected in a manner that minimizes statistical bias introduced to classical simulations caused by the MD practitioner “choosing” where simulations should start from. Each of these pre-selected seeds are run in parallel using short unbiased trajectories. Path exploration performed via adaptive sampling minimizes computing time while introducing greater control and efficiency towards observing rare/high energy protein structures from simulation.

Following the initial 100 ns production run, 50 states were selected to seed a new round of adaptive sampling. The seed states were selected using a least counts-based strategy, which helps identify the states that were least sampled during the simulation round (55). In this way, the provision of new structures for subsequent simulations introduces minimal statistical bias towards phase space exploration and the discovery of new protein structures. For monomer simulation trajectories, seed states were selected based off a vector containing the combined gating distances for Phe165_CB_– Phe43_CB_ and Arg70_CZ_–Asp190_OD2_ atoms.

Because a homotrimer was used, a sampling strategy had to be implemented which would strive to account for protomer-specific trimer dynamics. A vector combining the respective gating distances per protomer was calculated for each trimer trajectory. Then, the protomer-speciifc gating distances for the first trajectory frame, *n*, were compared to the next frame, *n+1*, using the Chebyshev distance. Whichever protomer’s gating distance experienced the maximum amount of conformational change from the preceding frame was used as the current frame’s placeholder in the output trajectory vector. This vector of Chebyshev-filtered gating distances was then provided for least counts-based clustering. All distances were calculated using MDTraj 1.9.3 (56).

Each round of clustering would yield a subsequent 50 trajectories from the 50 seed states for additional sampling. For monomer simulations, trajectories developed from each separate OsSWEET2b conformational state used in initial system construction (HG, IF, OC, and OF) underwent isolated rounds of adaptive sampling. Each monomer trajectory was run for 25 ns. Monomer systems beginning from different conformational states each underwent 11 rounds of adaptive sampling, totaling to 44 rounds of adaptive sampling per phosphorylation state (**Tables S4-7**).

OsSWEET2b trimer topologies began as inward-facing homotrimers and totaled ∼180,000 atoms, complicating adaptive sampling. To maximize data return and state exploration given computing wallclock times, each trimer seed state trajectory was run for 9 ns. Adaptive sampling proceeded until preliminary coverage of the HG, IF, OC, and OF states was observed. Then, 1000 seed states were selected based off least counts clustering of a combined vector list containing the protomer-specific gating distances calculated from all trimer trajectories. Each of these 1000 seed states were submitted to the Folding@Home distributed computing platform and run for 600 ns each (45, 46). This protocol was repeated for the trimer systems at each phosphorylation state.

Simulation frame save rates for all production simulations were recorded every 0.1 ns. Only trimer data obtained from Folding@Home runs is reported in this manuscript. Aggregate simulation amounted to: p_0_OsSWEET2b^M^ = 51.94 µs; p_2_OsSWEET2b^M^ = 55.58 µs; p_4_OsSWEET2b^M^ = 54.04 µs; p_0_OsSWEET2b^T^ = 588.62 µs; p_2_OsSWEET2b^T^ = 591.57 µs; and p_4_OsSWEET2b^T^ = 593.40 µs.

### Markov state model construction and validation

Markov state model (MSM) theory incorporates aggregate MD data into a master kinetic and thermodynamic framework (57–59). Briefly, all trajectories described by a set of molecular descriptors for a given system are provided as a list. This list can then be clustered into a finite number of microstates, creating a counts matrix. The counts matrix assigns each simulation frame to a cluster. Markov theory is then used to calculate the transition probability between states belonging to different clusters based off the counts. Describing simulation frames using feature sets which accurately depict rate-determining conformational change therefore becomes critical, as observed protein structures belonging to different microstate clusters should be geometrically and kinetically distinct (i.e., clustered are as homogeneous as possible).

The previously published OsSWEET2b feature set was used for featurization (24). The combined nine pairwise-atom-distance list of Asp190_CA_–Arg70_CA_, Asp190CA–Tyr62_CA_, Tyr184_CA_– Arg70_CA_, Phe165_CA_–Phe43_CA_, Glu164_CA_–Arg42_CA_, Ser139_OG_–Asn54_OD1_, Ser51_OG_–Gln84_OE1_, Leu170_CA_–Tyr48_CA_, and Ser171_CA_–Pro47_C_. Since these nine features have been previously used to describe OsSWEET2b conformational change (24, 26), these intraprotomer distance sets were calculated for each of trimer systems to generate a combined 27 distance list. After featurization, these feature sets underwent dimensionality reduction using time-lagged independent component analysis (tICA) (57). A connected tICA landscape, indicating reversible sampling throughout higher dimensional phase space, was required prior to MSM construction (**Figure S1**).

A grid search was conducted to optimize cluster count (100-1000 clusters, incremented by 50), tIC dimensionality (2-12 components), and Markov lag time hyperparameters for MSM construction. The tICA-reduced and clustered data was then used as MSM input. Markov lag times were first estimated using hyperparameter combinations from initial “dummy MSMs” where the implied timescales appeared converged. Grid search MSMs were evaluated based off their VAMP-2 score and the absolute difference between the eigenvalues of the second and third timescale dominant processes (t_2_-t_3_) (60, 61). Grid search MSMs which scored in the top 50 MSMs in maximizing both the VAMP-2 score and t_2_-t_3_ rankings had their implied timescale plots evaluated for convergence. The fastest time at which the implied timescale converged was selected as the official MSM lag time. The number of processes plotted for implied timescale analysis was determined via spectral decomposition analysis (62, 63) (**Figure S2**). The extent of stationary distribution reweighting was evaluated, with the goal of having the MSM-weighting deviate no more than 1.5 orders of magnitude above the raw distribution (**Figure S3**) (64). Lastly, the Chapman-Kolmogorov test was performed to evaluate detailed balance at Markov lag times greater than the validated lag time but smaller than the total trajectory length (**Figure S4**) (65–67).

All MSM-related calculations were made possible via Pyemma 2.5.6 (68). The final MSM hyperparameters are shown in **Table S8**.

### Bootstrapping analysis

MSM bootstrapping was performed across 200 data subsets, where each subset contained 80% of the total trajectories. After microstate clustering and MSM generation were completed for each subset, a binning protocol (24) was performed to project free energy errors with respect to gating landscapes over 100% of the data (**Figures S5 and S6**).

### Selection of representative trajectories for analysis

Representative trajectories were selected based off the identity of their metastable state. For OsSWEET2b monomer simulations, trajectories were chosen based off whether they existed entirely within an IF basin (extracellular gate: 0 < x < 5 Å; intracellular gate: 14 < y < 20 Å) or an OF basin (extracellular gate: 11 < x < 12 Å; intracellular gate: 5 < y < 6 Å). Because individual trimer trajectories were longer, protomer-specific gating states were more likely to change throughout each simulation. As such, IF-like trajectories (extracellular gate: 0 < x < 5 Å; intracellular gate: 14 < y < 20 Å) and OF-like trajectories (extracellular gate: 11 < x < 15 Å; intracellular gate: 5 < y < 10 Å) were determined so long as ≥ 30% of the trajectory frames resided within the aforementioned basins. Each selected trajectory was independently visualized atop the gating free energy landscapes to ensure that (1) the trajectory did indeed sample the specified state, and (2) the trajectory did not sample the opposing state at any time (i.e., a trajectory that was determined to be IF-like never transitioned to become OF-like). The number of trimer protomer states respectively presented in the IF-like, OF-like, or HG/OC-like conformations is summarized in **Figure S7**.

### Mean first passage time analysis

Mean first passage time (MFPT) was used to calculate the predicted rate for conformational transition between IF-to-OF states and OF-to-IF states for OsSWEET2b systems. Cluster identities for MFPT analyses were determined by seeing which clusters each of the frames from the representative trajectories belonged. The top five resulting clusters which constituted at least 5% of all reported clusters for the metastable state were then used with proceeding calculations. For trimer simulations, MFPT was calculated in a protomer-dependent fashion, meaning that clusters were identified for the IF-to-OF and OF-to-IF transitions observed per protomer. Using the top five clusters as described above, MFPT calculations were performed upon each of the bootstrapped MSM to enable statistical confidence. Outliers were removed based off the 1.5x-interquartile range rule. Statistical significance was then performed using Welch’s t-test. All MFPT statistics are reported in **Table S2.**

### Hydrogen bonding analysis

The CPPTRAJ hbond command was used to calculate hydrogen bonds between any atoms between the C-terminal tail residues and the remainder of the protein. For trimer structures, only intraprotomer hydrogen bonds were calculated. This method first led to identify which residues participated in C-terminal tail hydrogen bonds using the representative trajectories. Redundancy in calculations was removed by only accounting for the residue functional groups which belonged to the greatest number of hydrogen bond-participating frames. Analysis was focused on intraprotomer C-terminal – cytosolic loop hydrogen bonding interactions. Filtered hydrogen bonding frequencies were summed across all representative trajectories and are reported in **Tables S3 and S4**. CPPTRAJ v5.1.0 was used (69).

### Trajectory handling

CPPTRAJ v5.1.0 was used for all trajectory, coordinate, and topology file processing and conversion necessary for analyses and simulations.

### Data visualization and representation

Trajectories were visualized using VMD (70). PDB structures were visualized using ChimeraX (71). Plots were generated using Matplotlib 3.0.3 and Plotly 5.13.0 using Python 3.6. Plots were visualized using Jupyter Notebooks. Final figures were prepared using Adobe Illustrator 2021.

## DATA AVAILABILITY

We currently make our data available upon request. We intend to prepare a public repository of codes and data upon final publication.

## SUPPORTING INFORMATION

This article contains supporting information.

## ACKNOWLEDGEMENTS

The authors thank Prof. Li-Qing Chen (University of Illinois, Urbana Champaign) for recommendation of bioinformatic tools, specifically GPS, for probing putative phosphorylation patterns, as well as review of earlier drafts of this manuscript.

## Funding and additional information

This work has been supported by the National Institutes of Health (Award No. R35GM142745). Molecular dynamics simulations of monomeric OsSWEET2b were performed on the Blue Waters Supercomputer at the National Center for Supercomputing Applications (NCSA), as well as the NCSA Illinois Delta Early Access Initiative. The Delta advanced computing and data resource is supported by the National Science Foundation (award OAC 2005572) and the State of Illinois. Delta, as well as Blue Waters during its tenure, exist(ed) as a joint effort for the University of Illinois Urbana-Champaign and its NCSA. Production runs of trimeric OsSWEET2b were performed using the Folding@Home distributed computing platform (Project IDs 17916, 17917, and 17918). The authors give special thanks to the individual internal testers, beta testers, and folders that have helped make this work possible.

