## Supplementary Information for "Interplay between phosphorylation and oligomerization tunes the conformational ensemble of SWEET transporters"

### Table of Contents

#### Figures and Tables

Table S1 Bioinformatic prediction of OsSWEET2b phosphorylation site patterning S-3

Table S2 Transition rates for OsSWEET2b proteoform conformational changes associated sugar export or import S-4

Table S3 Hydrogen bonding (hbond) analysis between monomer C-terminal tail and cytosolic loops S-5

Table S4 Intraprotomer hydrogen bonding (hbond) analysis between trimer C-terminal tail and cytosolic loops S-6

Table S5 $p_0$OsSWEET2b$^M$ adaptive sampling regime S-8

Table S6 $p_2$OsSWEET2b$^M$ adaptive sampling regime S-9

Table S7 $p_4$OsSWEET2b$^M$ adaptive sampling regime S-10

Table S8 Markov state model (MSM) hyperparameter selection S-11

Figure S1 time-lagged independent component analysis (tICA) free energy landscapes for OsSWEET2b monomer and trimer phosphorylation state systems S-12

Figure S2 Implied timescale plots for OsSWEET2b monomer and trimer phosphorylation state systems S-13

Figure S3 Raw counts versus MSM population for each microstate cluster of OsSWEET2b monomer and trimer phosphorylation state systems S-14

Figure S4 Chapman-Kolmogorov tests for final OsSWEET2b MSMs S-15

Figure S5 Gating free energy error estimates on $p_n$OsSWEET2b$^M$ systems from 200 bootstrapped Markov state models S-16

Figure S6 Gating free energy error estimates on $p_n$OsSWEET2b$^T$ systems from 200 bootstrapped Markov state models S-17

Figure S7 Bullet indicator plots representing protomer conformational diversity among representative OsSWEET2b$^T$ trajectories S-18

### SUPPORTING INFORMATION

**Table S1. Bioinformatic prediction of OsSWEET2b phosphorylation site patterning.** TK = tyrosine kinase; TKL = tyrosine kinase-like; INSR = insulin receptor tyrosine kinase; PKC = protein kinase c (serine/threonine kinase); PKA = protein kinase a (serine/threonine kinase)

| OsSWEET2b residue | Software | Score | Suggested kinases |
|---|---|---|---|
| **Tyr214** | GPS-6.0 | 0.962 / 1.000 | TK |
| | NetPhos-3.1 | 0.367 / 1.000 | INSR |
| **Ser215** | GPS-6.0 | 0.070 / 1.000 | TKL |
| | NetPhos-3.1 | 0.847 / 1.000 | PKC |
| **Ser223** | GPS-6.0 | 0.291 / 1.000 | TKL |
| | NetPhos-3.1 | 0.518 / 1.000 | PKA |
| | MusiteDeep | 0.518 / 1.000 | N/A |
| **Ser224** | GPS-6.0 | 0.331 / 1.000 | TKL |
| | NetPhos-3.1 | 0.455 / 1.000 | PKA |
| | MusiteDeep | 0.560 / 1.000 | N/A |

**Table S2. Transition rates for OsSWEET2b proteoform conformational changes associated sugar export or import.** *Apo* export transitions are characterized by transition from an IF state to an OF state. *Apo* import transitions are characterized by transition from an OF state to an IF state. Statistical significance evaluated using Welch's t-test after removal of outliers. Standard deviations reported (S.D.). Monomer sample sizes: $p_0$OsSWEET2b$^{M;IF\text{-}to\text{-}OF}$, n=181; $p_0$OsSWEET2b$^{M,OF\text{-}to\text{-}IF}$, n=1931; $p_2$OsSWEET2b$^{M;IF\text{-}to\text{-}OF}$, n=192; $p_2$OsSWEET2b$^{M;OF\text{-}to\text{-}IF}$, n=196; $p_4$OsSWEET2b$^{M;IF\text{-}to\text{-}OF}$, n=194; $p_4$OsSWEET2b$^{M;OF\text{-}to\text{-}IF}$, n=193. Trimer sample sizes: $p_0$OsSWEET2b$^{T,A;IF\text{-}to\text{-}OF}$, n=186; $p_0$OsSWEET2b$^{T,A\text{-}import}$, n=194; $p_0$OsSWEET2b$^{T,B;IF\text{-}to\text{-}OF}$, n=186; $p_0$OsSWEET2b$^{T,B\text{-}import}$, n=180; $p_0$OsSWEET2b$^{T,C;\ IF\text{-}to\text{-}OF}$, n=191; $p_0$OsSWEET2b$^{T,C\text{-}import}$, n=195;
$p_2$OsSWEET2b$^{T,A;IF\text{-}to\text{-}OF}$, n=193; $p_2$OsSWEET2b$^{T,A\text{-}import}$, n=197; $p_2$OsSWEET2b$^{T,B;IF\text{-}to\text{-}OF}$, n=189; $p_2$OsSWEET2b$^{T,B\text{-}import}$, n=196; $p_2$OsSWEET2b$^{T,C;IF\text{-}to\text{-}OF}$, n=194; $p_2$OsSWEET2b$^{T,C\text{-}import}$, n=191
$p_4$OsSWEET2b$^{T,A;IF\text{-}to\text{-}OF}$, n=200; $p_4$OsSWEET2b$^{T,A\text{-}import}$, n=194; $p_4$OsSWEET2b$^{T,B;IF\text{-}to\text{-}OF}$, n=198; $p_4$OsSWEET2b$^{T,B\text{-}import}$, n=195; $p_4$OsSWEET2b$^{T,C;IF\text{-}to\text{-}OF}$, n=196; $p_4$OsSWEET2b$^{T,C\text{-}import}$, n=197.

| Monomer Proteoform | Mean IF-to-OF transition rate (μs) | S.D. IF-to-OF transition | Mean OF-to-IF transition rate (μs) | S.D. OF-to-IF transition | p-value |
|---|---|---|---|---|---|
| **$p_0$OsSWEET2b$^M$** | 3.0789 | 1.3474 | 3.5287 | 1.500 | 2.50e-03 |
| **$p_2$OsSWEET2b$^M$** | 1.5297 | 0.3841 | 8.3374 | 4.5041 | 2.74e-64 |
| **$p_4$OsSWEET2b$^M$** | 1.2943 | 0.4406 | 2.2044 | 0.6031 | 1.45e-22 |
| **Trimer Proteoform** | **Mean IF-to-OF transition rate (μs)** | **S.D. IF-to-OF transition** | **Mean OF-to-IF transition rate (μs)** | **S.D. OF-to-IF transition** | **p-value** |
| **$p_0$OsSWEET2b$^{T,A}$** | 13.9249 | 3.0366 | 23.7874 | 7.2673 | 4.32e-49 |
| **$p_0$OsSWEET2b$^{T,B}$** | 13.9249 | 3.0366 | 18.2399 | 5.6263 | 1.52e-64 |
| **$p_0$OsSWEET2b$^{T,C}$** | 5.7220 | 0.8612 | 47.2105 | 8.9743 | 1.45e-93 |
| **$p_2$OsSWEET2b$^{T,A}$** | 6.9382 | 0.9414 | 35.7089 | 7.6552 | 1.78e-176 |
| **$p_2$OsSWEET2b$^{T,B}$** | 8.8961 | 1.3742 | 25.5665 | 5.6100 | 2.46e-206 |
| **$p_2$OsSWEET2b$^{T,C}$** | 8.7024 | 1.2930 | 28.6888 | 8.5008 | 5.03e-151 |
| **$p_4$OsSWEET2b$^{T,A}$** | 3.3189 | 0.2826 | 73.4235 | 13.9493 | 5.02e-226 |
| **$p_4$OsSWEET2b$^{T,B}$** | 97.8033 | 25.1083 | 14.9475 | 3.3371 | 3.96e-34 |
| **$p_4$OsSWEET2b$^{T,C}$** | 128.4513 | 26.4650 | 7.7166 | 1.0660 | 4.01e-100 |

**Table S3. Hydrogen bonding (hbond) analysis between monomer C-terminal tail and cytosolic loops.** Hbond analysis was performed on representative trajectories that primarily sampled either an IF-like or an OF-like state.

| OsSWEET2b Protomer | hbond | Abundance (%) | No. of frames | Total frames |
|---|---|---|---|---|
| **$p_0$OsSWEET2b$^{M\text{-}IF}$** | Ser224-Val163 | 8.2784 | 515 | 6221 |
| | Ser224-Glu164 | 5.6422 | 351 | 6221 |
| **$p_0$OsSWEET2b$^{M\text{-}OF}$** | Tyr214-Ser160 | 9.4567 | 644 | 6810 |
| | Arg219-Glu161 | 10.8370 | 738 | 6810 |
| | Asp222-Glu161 | 6.7401 | 459 | 6810 |
| | Ser223-Glu161 | 12.4229 | 846 | 6810 |
| | Ser224-Glu161 | 14.8899 | 1014 | 6810 |
| **$p_2$OsSWEET2b$^{M\text{-}IF}$** | Tyr214-Glu161 | 7.8400 | 588 | 7500 |
| | Tyr214-Val163 | 15.2667 | 1145 | 7500 |
| | Trp218-Val163 | 6.2800 | 471 | 7500 |
| **$p_2$OsSWEET2b$^{M\text{-}OF}$** | Ser215-Arg159 | 7.1143 | 249 | 3500 |
| | Ser224-Arg159 | 47.0857 | 1648 | 3500 |
| | Ser215-Ser162 | 13.4857 | 472 | 3500 |
| **$p_4$OsSWEET2b$^{M\text{-}IF}$** | — | — | — | 2750 |
| **$p_4$OsSWEET2b$^{M\text{-}OF}$** | Leu229-Ser39 | 7.5300 | 614 | 8154 |
| | Tyr214-Ser162 | 24.9080 | 2031 | 8154 |

**Table S4. Intraprotomer hydrogen bonding (hbond) analysis between trimer C-terminal tail and cytosolic loops.** Hbond analysis was performed on representative trajectories that primarily sampled either an IF-like or an OF-like state.

| OsSWEET2b Protomer | hbond | Abundance (%) | No. of frames | Total frames |
|---|---|---|---|---|
| $p_0OsSWEET2b^{T,A-IF}$ | Ser215-Glu164 | 13.5179 | 7301 | 54010 |
| $p_0OsSWEET2b^{T,B-IF}$ | — | — | — | 12000 |
| $p_0OsSWEET2b^{T,C-IF}$ | — | — | — | 18000 |
| $p_0OsSWEET2b^{T,A-OF}$ | Tyr214-Glu161 | 6.9636 | 12490 | 179362 |
| $p_0OsSWEET2b^{T,B-OF}$ | — | — | — | 198000 |
| $p_0OsSWEET2b^{T,C-OF}$ | Trp218-Glu161 | 6.9382 | 9991 | 144000 |
| $p_2OsSWEET2b^{T,A-IF}$ | — | — | — | 78000 |
| $p_2OsSWEET2b^{T,B-IF}$ | Ser224-Arg42 | 29.4375 | 942 | 3200 |
| | Pro226-Arg42 | 11.7188 | 375 | 3200 |
| | Leu227-Arg42 | 10.4063 | 333 | 3200 |
| | Leu228-Arg159 | 5.5000 | 176 | 3200 |
| | Tyr214-Glu161 | 11.6563 | 373 | 3200 |
| | Ser224-Ser162 | 12.8750 | 412 | 3200 |
| | Ala225-Ser162 | 5.2500 | 168 | 3200 |
| | Arg219-Val163 | 63.4375 | 2030 | 3200 |
| | Tyr214-Glu164 | 60.0313 | 1921 | 3200 |
| | Arg219-Glu164 | 35.1250 | 1124 | 3200 |
| | Ser223-Glu164 | 20.8750 | 668 | 3200 |
| | Pro226-Glu164 | 11.6875 | 374 | 3200 |
| $p_2OsSWEET2b^{T,C-IF}$ | — | — | — | 18000 |
| $p_2OsSWEET2b^{T,A-OF}$ | Arg219-Glu161 | 10.3085 | 14102 | 136800 |
| $p_2OsSWEET2b^{T,B-OF}$ | — | — | — | 311962 |
| $p_2OsSWEET2b^{T,C-OF}$ | — | — | — | 315182 |
| $p_4OsSWEET2b^{T,A-IF}$ | Ser223-Arg42 | 31.8075 | 76931 | 241864 |
| | Ser223-Arg159 | 14.2175 | 34387 | 241864 |
| | Ser223-Ser162 | 5.3514 | 12943 | 241864 |
| | Asp222-Val163 | 5.4423 | 13163 | 241864 |
| | Arg219-Glu164 | 5.4522 | 13187 | 241864 |
| $p_4OsSWEET2b^{T,B-IF}$ | Leu229-Lys38 | 10.7917 | 2590 | 24000 |
| | Ser223-Arg42 | 10.1500 | 2436 | 24000 |
| | Arg219-Glu164 | 6.8333 | 1640 | 24000 |
| $p_4OsSWEET2b^{T,C-IF}$ | Ser215-Ser162 | 7.3368 | 10565 | 144000 |
| $p_4OsSWEET2b^{T,A-OF}$ | Ser223-Lys38 | 7.3491 | 9587 | 130452 |
| | Ser223-Arg42 | 37.4866 | 48902 | 130452 |
| | Ser224-Arg42 | 5.7699 | 7527 | 130452 |
| | Ser223-Ser162 | 6.3702 | 8310 | 130452 |

| | | | | |
|---|---|---|---|---|
| | Arg219-Glu164 | 7.3652 | 9608 | 130452 |
| **$p_4$OsSWEET2b$^{T,B\text{-}OF}$** | Asp222-Lys38 | 11.7127 | 25960 | 221640 |
| | Arg219-Ser162 | 13.7692 | 30518 | 221640 |
| | Gly220-Val163 | 8.7186 | 19324 | 221640 |
| | Gly220-Glu164 | 25.0546 | 55531 | 221640 |
| | Gln221-Glu164 | 13.2891 | 29454 | 221640 |
| **$p_4$OsSWEET2b$^{T,C\text{-}OF}$** | Asp222-Arg42 | 5.8778 | 1058 | 18000 |
| | Ser223-Arg159 | 22.8333 | 4110 | 18000 |
| | Arg219-Glu161 | 7.4944 | 1349 | 18000 |
| | Asp222-Ser162 | 14.6333 | 2634 | 18000 |

**Table S5. $p_0$OsSWEET2b$^M$ adaptive sampling regime.**

| $p_0$OsSWEET2b$^M$ Round | Parallel Simulations (Trajectories) | | | | Target Trajectory Length (ns) | Round Simulation Time (µs) |
|---|---|---|---|---|---|---|
| | HG (start) | IF (start) | OC (start) | OF (start) | | |
| **0** | 1 | 1 | 1 | 1 | 100 | 0.35 |
| **1** | 50 | 50 | 50 | 50 | 250 | 4.14 |
| **2** | 50 | 50 | 50 | 50 | 250 | 4.79 |
| **3** | 50 | 50 | 50 | 50 | 250 | 4.75 |
| **4** | 50 | 50 | 50 | 50 | 250 | 5.00 |
| **5** | 50 | 50 | 50 | 50 | 250 | 4.98 |
| **6** | 50 | 50 | 50 | 50 | 250 | 4.13 |
| **7** | 50 | 50 | 39 | 32 | 250 | 4.23 |
| **8** | 50 | 50 | 50 | 50 | 250 | 4.96 |
| **9** | 50 | 50 | 50 | 50 | 250 | 5.00 |
| **10** | 50 | 50 | 50 | 50 | 250 | 4.61 |
| **11** | 50 | 50 | 50 | 50 | 250 | 4.99 |
| **Total Simulation Time**: *51.94 µs* | | | | | | |

Table S6. $p_2$OsSWEET2b$^M$ adaptive sampling regime.

| $p_2$OsSWEET2b$^M$ Round | Parallel Simulations (Trajectories) | | | | Target Trajectory Length (ns) | Round Simulation Time (µs) |
|---|---|---|---|---|---|---|
| | HG (start) | IF (start) | OC (start) | OF (start) | | |
| **0** | 1 | 1 | 1 | 1 | 100 | 0.45 |
| **1** | 52 | 49 | 52 | 52 | 250 | 5.16 |
| **2** | 49 | 50 | 50 | 50 | 250 | 4.98 |
| **3** | 50 | 50 | 50 | 50 | 250 | 5.00 |
| **4** | 50 | 50 | 50 | 50 | 250 | 5.00 |
| **5** | 50 | 50 | 50 | 50 | 250 | 5.00 |
| **6** | 50 | 50 | 50 | 50 | 250 | 5.00 |
| **7** | 50 | 50 | 39 | 32 | 250 | 5.00 |
| **8** | 50 | 50 | 50 | 50 | 250 | 5.00 |
| **9** | 50 | 50 | 50 | 50 | 250 | 5.00 |
| **10** | 50 | 50 | 50 | 50 | 250 | 5.00 |
| **11** | 50 | 50 | 50 | 50 | 250 | 5.00 |
| **Total Simulation Time**: *55.58 µs* | | | | | | |

**Table S7. $p_4$OsSWEET2b$^M$ adaptive sampling regime.**

| $p_4$OsSWEET2b$^M$ Round | Parallel Simulations (Trajectories) | | | | Target Trajectory Length (ns) | Round Simulation Time (µs) |
|---|---|---|---|---|---|---|
| | HG (start) | IF (start) | OC (start) | OF (start) | | |
| **0** | 1 | 1 | 1 | 1 | 100 | 0.36 |
| **1** | 50 | 50 | 50 | 50 | 250 | 4.91 |
| **2** | 50 | 50 | 50 | 50 | 250 | 4.81 |
| **3** | 50 | 50 | 50 | 50 | 250 | 4.99 |
| **4** | 50 | 50 | 50 | 50 | 250 | 4.90 |
| **5** | 50 | 50 | 50 | 50 | 250 | 4.99 |
| **6** | 50 | 50 | 50 | 50 | 250 | 4.11 |
| **7** | 50 | 50 | 50 | 50 | 250 | 5.00 |
| **8** | 50 | 50 | 50 | 50 | 250 | 4.97 |
| **9** | 50 | 50 | 50 | 50 | 250 | 5.00 |
| **10** | 50 | 50 | 50 | 50 | 250 | 5.00 |
| **11** | 50 | 50 | 50 | 50 | 250 | 5.00 |
| **Total Simulation Time**: *54.04 µs* | | | | | | |

**Table S8. Markov state model (MSM) hyperparameter selection**

| Proteoform | Cluster count | tICs | MSM lag time (ns) |
|---|---|---|---|
| **$p_0$OsSWEET2b$^M$** | 750 | 5 | 15 |
| **$p_2$OsSWEET2b$^M$** | 300 | 4 | 15 |
| **$p_4$OsSWEET2b$^M$** | 300 | 5 | 15 |
| **$p_0$OsSWEET2b$^T$** | 200 | 5 | 250 |
| **$p_2$OsSWEET2b$^T$** | 200 | 7 | 250 |
| **$p_4$OsSWEET2b$^T$** | 100 | 3 | 250 |

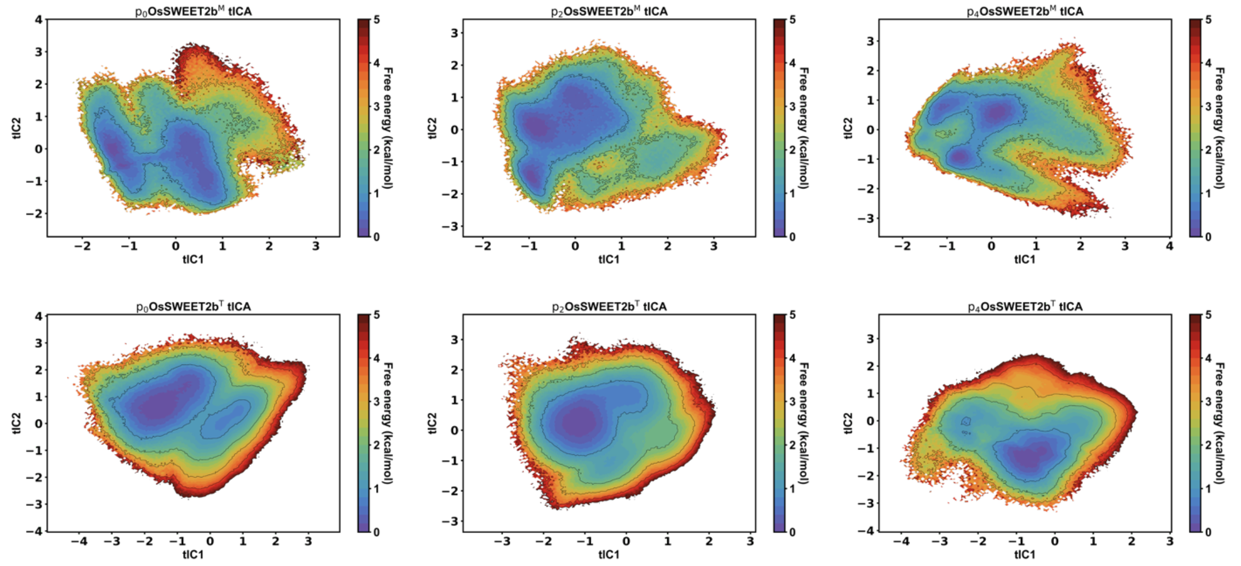

**Figure S1. time-lagged independent component analysis (tICA) free energy landscapes for OsSWEET2b monomer and trimer phosphorylation state systems.** These tICA decompositions were used as inputs for Markov state model construction. Free energy is reported in kcal/mol.

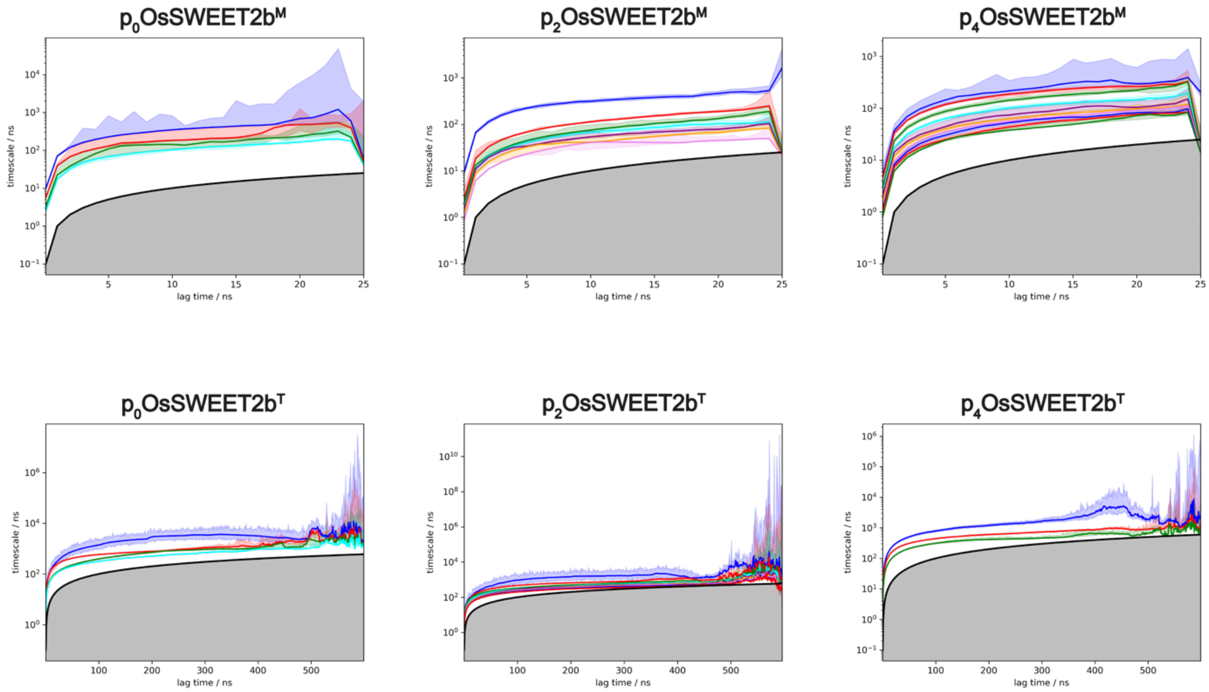

**Figure S2. Implied timescale plots for OsSWEET2b monomer and trimer phosphorylation state systems.**

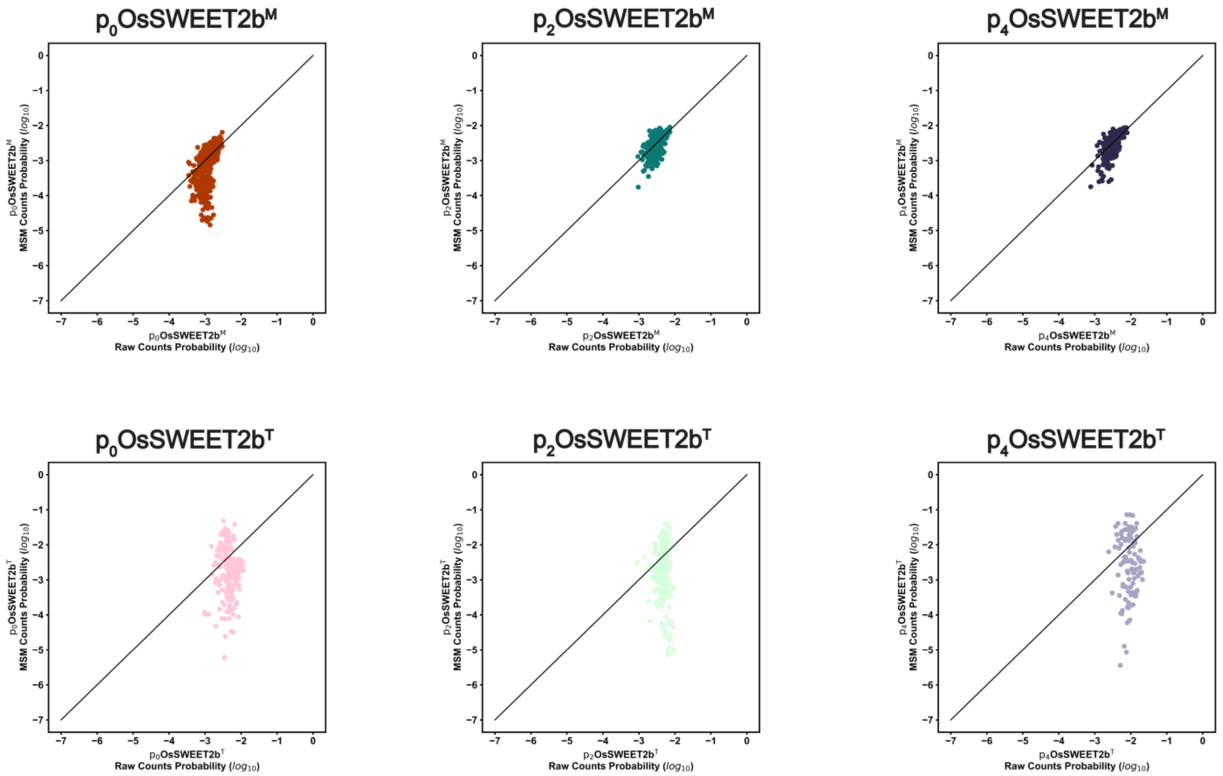

**Figure S3. Raw counts versus MSM population for each microstate cluster of OsSWEET2b monomer and trimer phosphorylation state systems.**

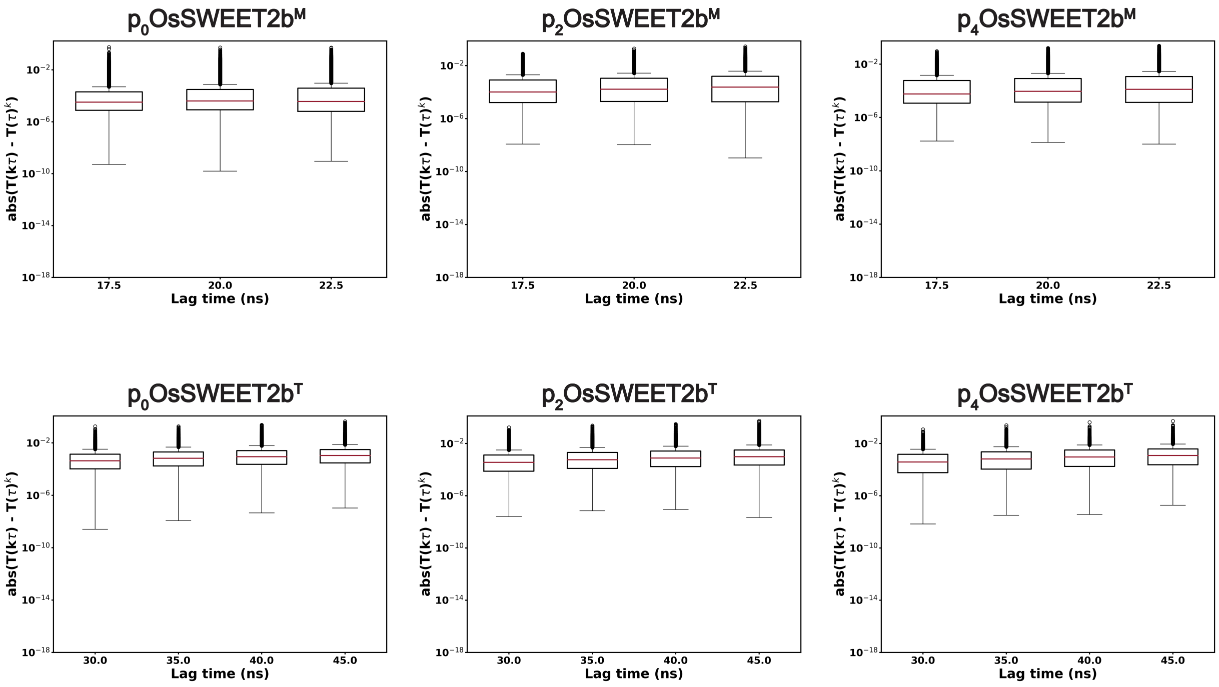

**Figure S4. Chapman-Kolmogorov tests for final OsSWEET2b MSMs.**

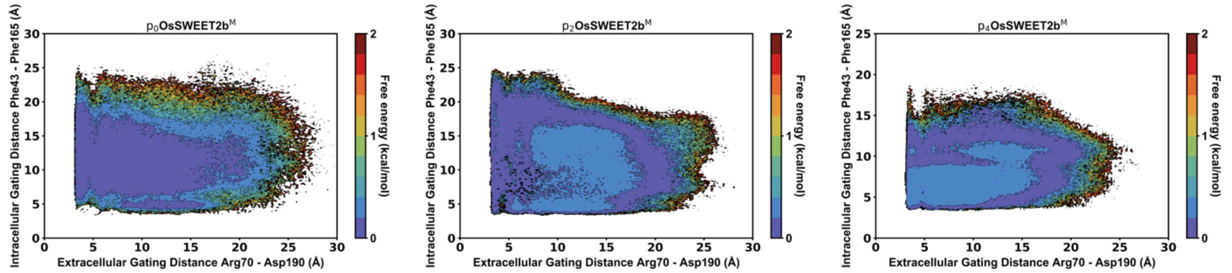

**Figure S5. Gating free energy error estimates on $p_n$OsSWEET2b$^M$ systems from 200 bootstrapped Markov state models.** Error calculations performed for monomer systems with no (*left*), $P_2$ (*middle*), and $P_4$ (*right*) phosphorylation patterns. Gating distances are plotted in angstroms (Å) with free energy reported in kcal/mol.

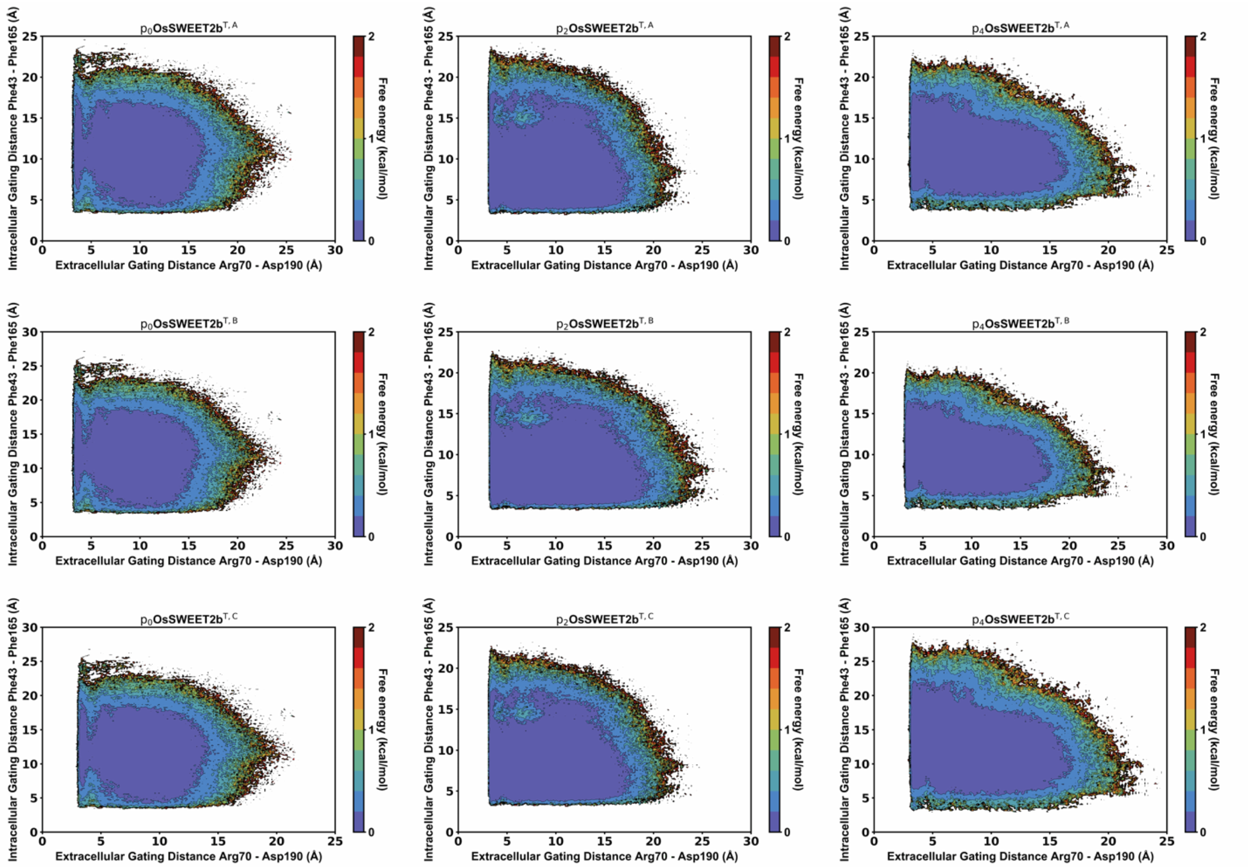

**Figure S6. Gating free energy error estimates on $p_n$OsSWEET2b$^T$ systems from 200 bootstrapped Markov state models.** Error calculations performed for trimer systems with no (*left column*), $P_2$ (*middle column*), and $P_4$ (*right column*) phosphorylation patterns. Individual protomers are shown for chains A (*top row*), B (*middle row*), and C (*bottom row*). Gating distances are plotted in angstroms (Å) with free energy reported in kcal/mol.

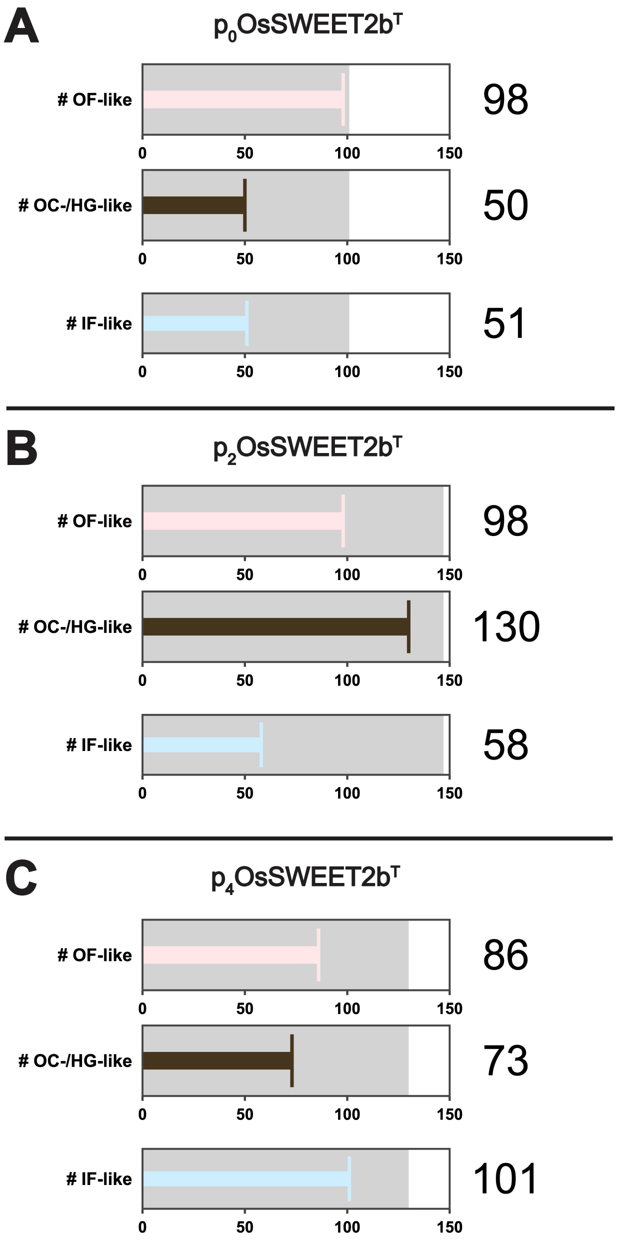

**Figure S7. Bullet indicator plots representing protomer conformational diversity among representative OsSWEET2b$^T$ trajectories.** Classification and counts of different metastable conformations seen from individual trajectories are shown for (**A**) $p_0$OsSWEET2b$^T$, (**B**) $p_2$OsSWEET2b$^T$, and (**C**) $p_0$OsSWEET2b$^T$. The "bullet" indicator depicts how many protomers were observed for a given state within all the trajectories. The grey background indicates the total number of trajectories within the "representative" pool used for gating timeseries and C-terminal hydrogen bonding analyses.
